## Supplementary Figures and Tables 1 and 8 for "SMCHD1 has separable roles in chromatin architecture and gene silencing that could be targeted in disease"

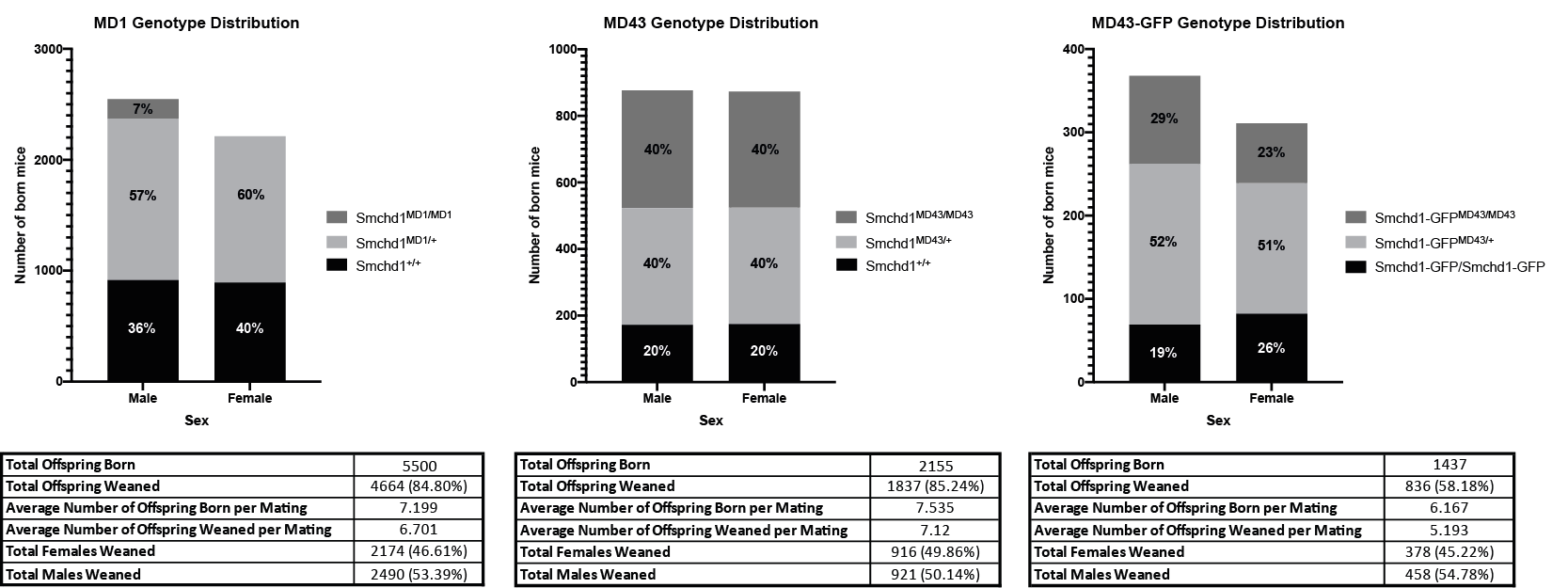

**Supplementary Figure 1.** Summarised genotype distribution data at weaning for offspring from for heterozygous intercrosses for *MommeD1*, *MommeD43* and *GFP-MommeD43*. *MommeD43* abbreviated to *MD43*. *MommeD1* abbreviated to *MD1.*

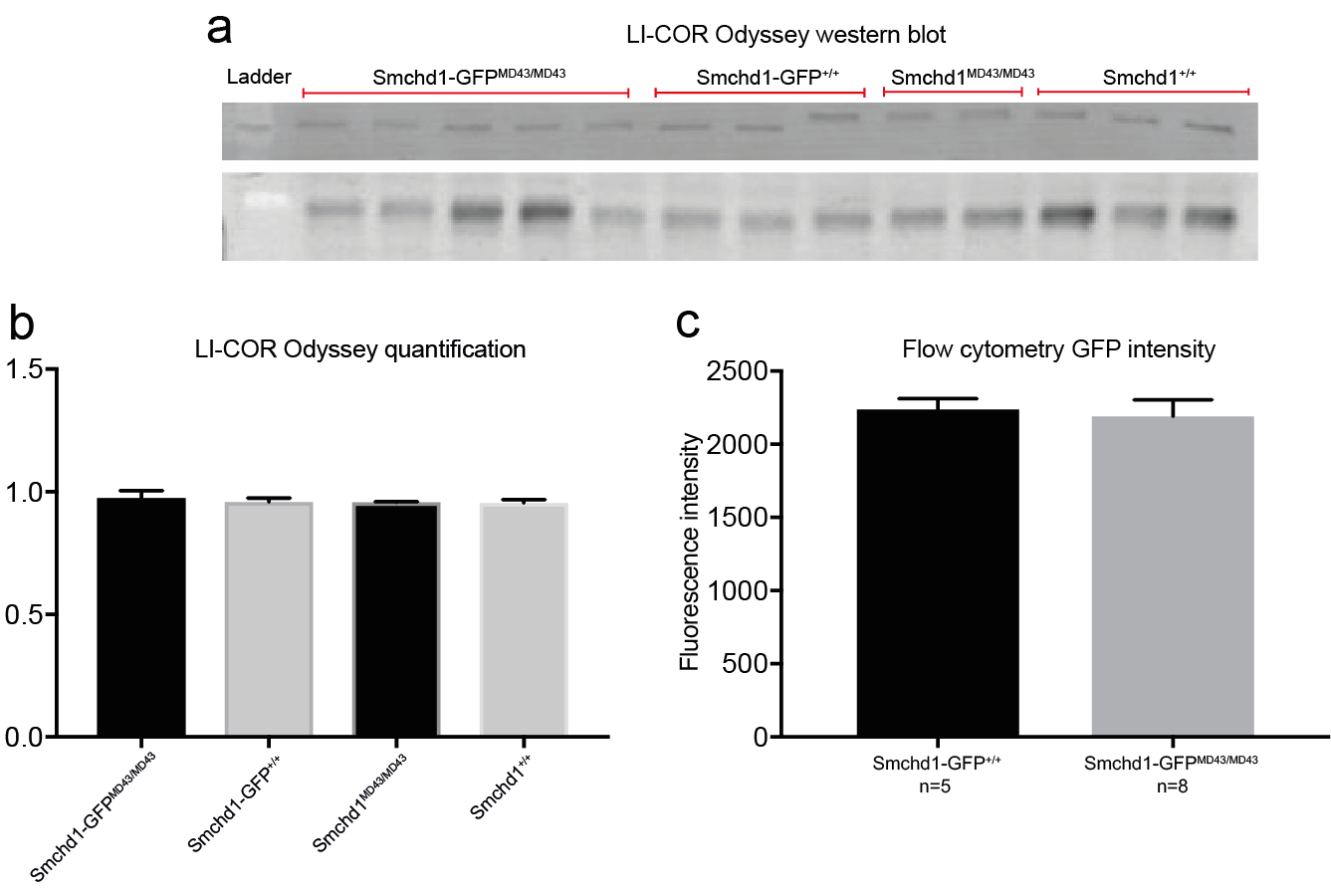

**Supplementary Figure 2.** a. LI-COR Odyssey quantitative western blot picture to measure levels of SMCHD1 protein in *Smchd1^GFP-MommeD43/GFP-MommeD43^*, *Smchd1^GFP/GFP^*, *Smchd1^MommeD43/MommeD43^* and *Smchd1^+/+^* primary MEFs (side-by-side lanes of the same genotype are biological replicates). b. Quantification of the LI-COR Odyssey western blot shown in a. c. Mean GFP fluorescence intensity in *Smchd1^GFP-MommeD43/GFP-MommeD43^*and *Smchd1^GFP/GFP^* primary MEFs measured by flow cytometry showing the same levels of SMCHD1 protein. Number of biological replicates indicated below. *MommeD43* abbreviated to *MD43*.

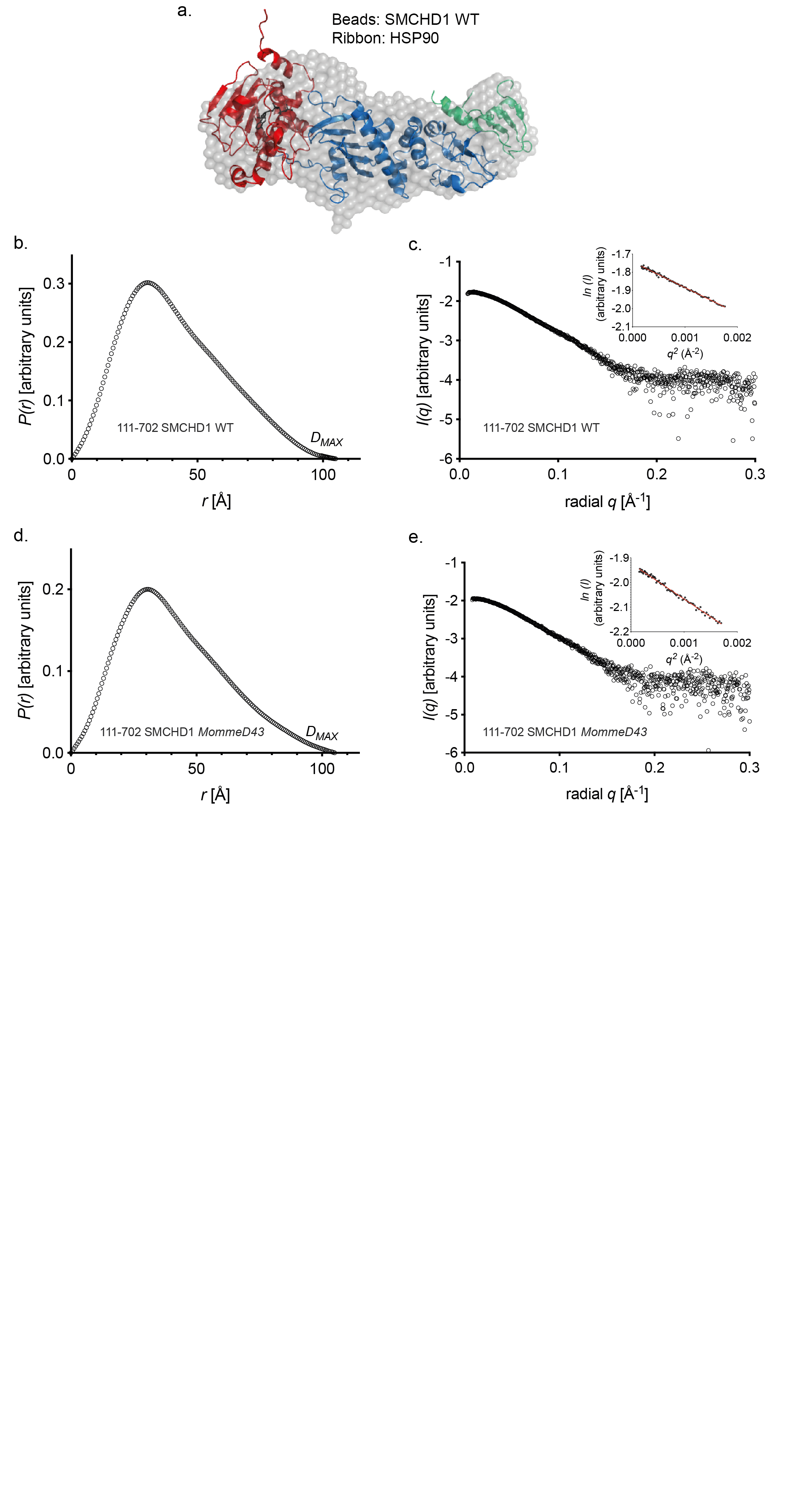

**Supplementary Figure 3**: a. Bead model of the extended ATPase domain (aa 111-702) of murine wild-type SMCHD1 (SMCHD1 *MommeD43* has the same structure). Overlayed is the ribbon structure of the highly related GHKL ATPase domain of HSP90. b., d. Distance distribution function ( P(r) ) calculated from data shown in c. for wild-type Smchd1 and e. for SMCHD1 MommeD43. c., e. Experimental scattering curves (black circles) of the recombinant ATPase domain of SMCHD1 WT and SMCHD1 MommeD43 respectively. On the right upper corner are the corresponding Guinier plots with the trend line from linear regression in red.

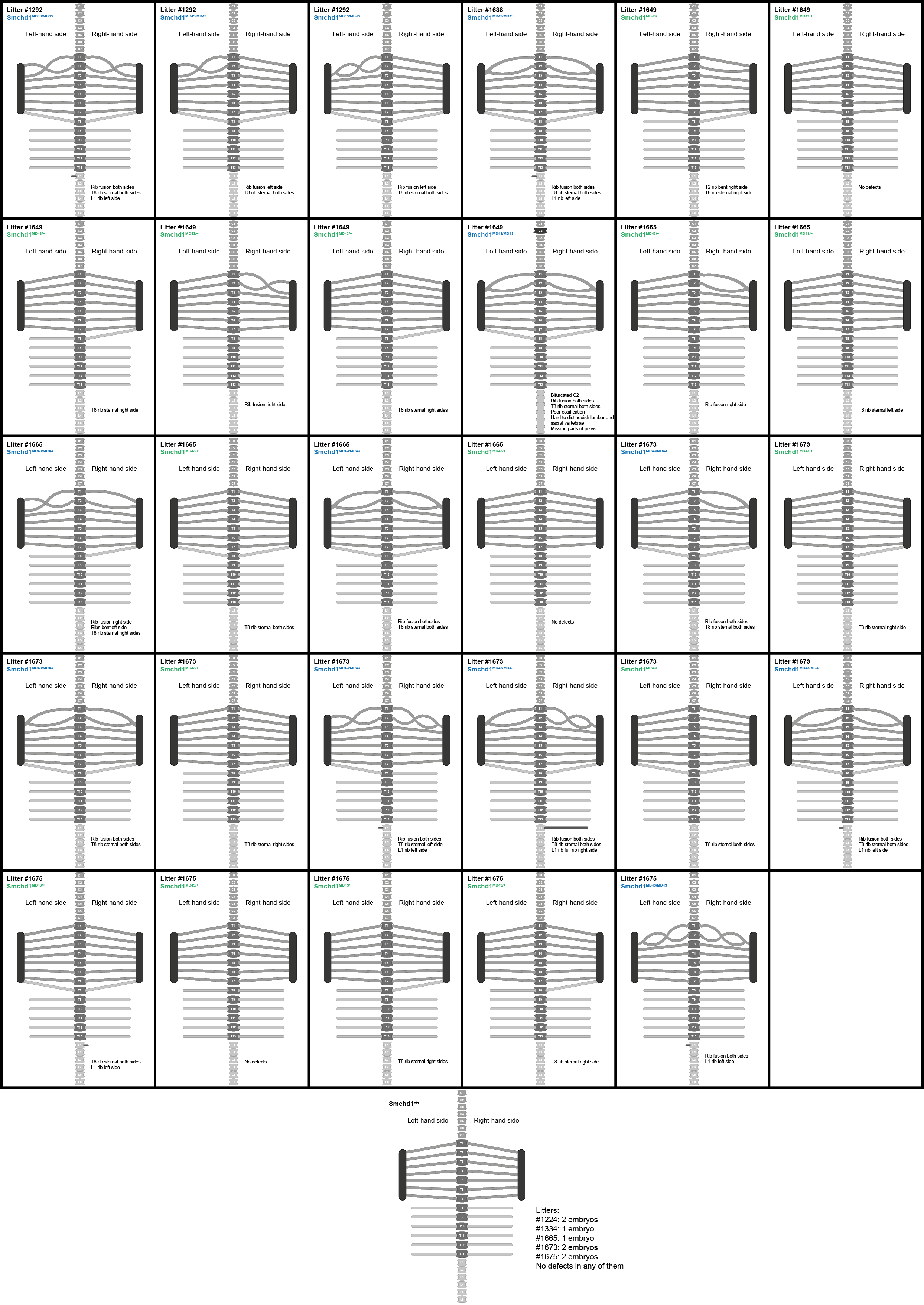

**Supplementary Figure 4.** Detailed scoring of skeletal defects observed in *Smchd1^MommeD43/MommeD43^*, *Smchd1^MommeD43/+^* and *Smchd1^+/+^* E16.5 littermate embryos**.** Schematic representation of each embryo skeleton scored after skeletal preparation (37 in total). Only one diagram at the bottom represents all 8 Smchd1^+/+^ embryos since all of them are identical. For the others, approximations of the shapes of the observed fused ribs were drawn. The view is dorsal and the sternum is represented on both sides by the dark vertical bar to which the T1 to T7 ribs join in the wild-type embryos. *MommeD43* abbreviated to *MD43*.

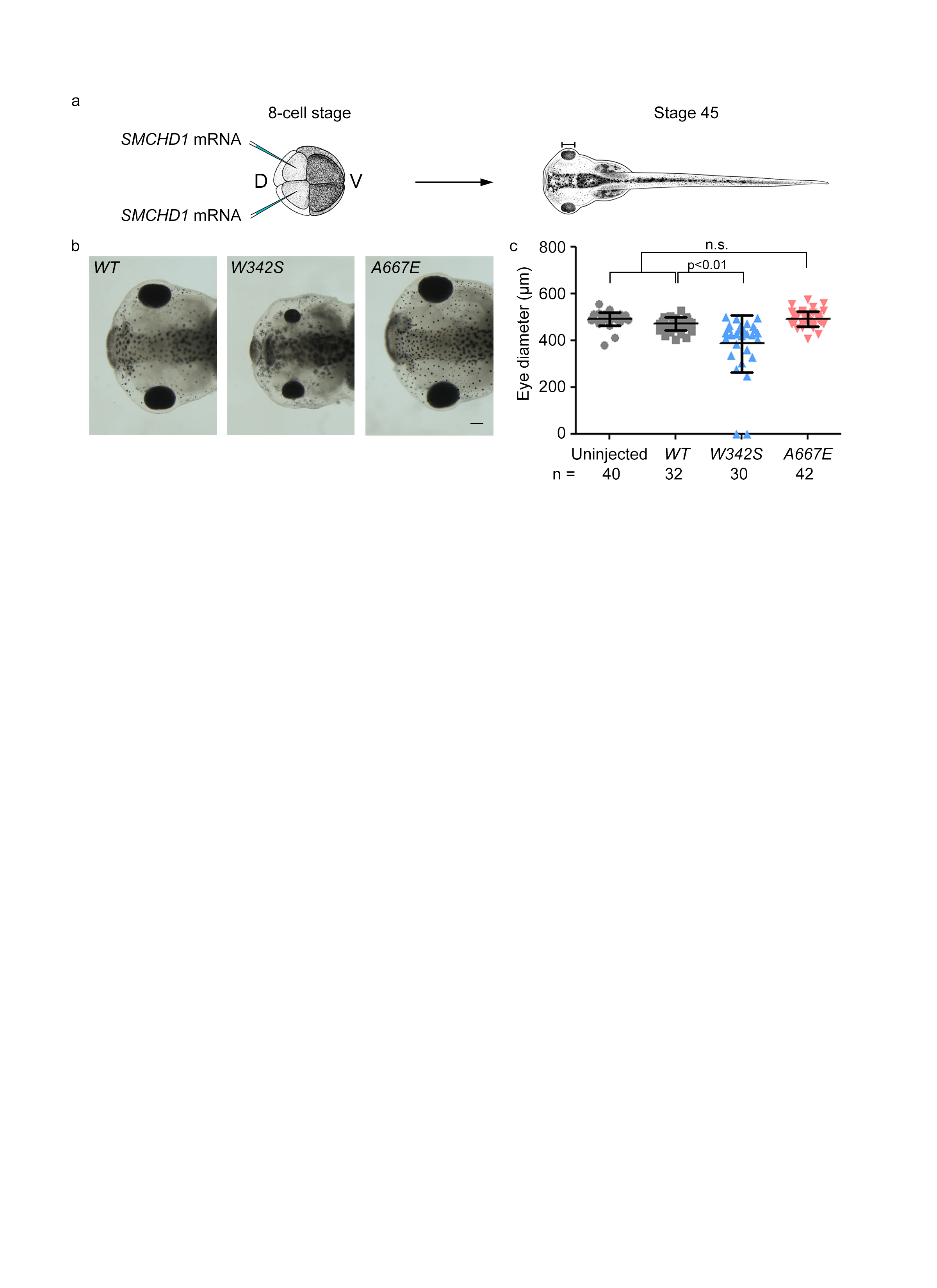

**Supplementary Figure 5.** Overexpression of *MommeD43* (A667E) mutation in *Xenopus laevis* embryos does not cause craniofacial anomalies. (a) Schematic of experiment. *SMCHD1* mRNA was injected into the two dorsal (D) animal blastomeres at the 8-cell stage to target the future head of the tadpole. Eye diameters of the tadpoles were observed at Stage 45. (b) Representative pictures of tadpoles injected with *SMCHD1* carrying various mutations. Scale bar represent 200 μm. (c) Measurements of eye diameters in tadpoles injected with various SMCHD1 mRNA. W342S is a BAMS mutation, A667E is the *MommeD43* mutation. Results are given as means ± standard deviation. n.s.: not significant.

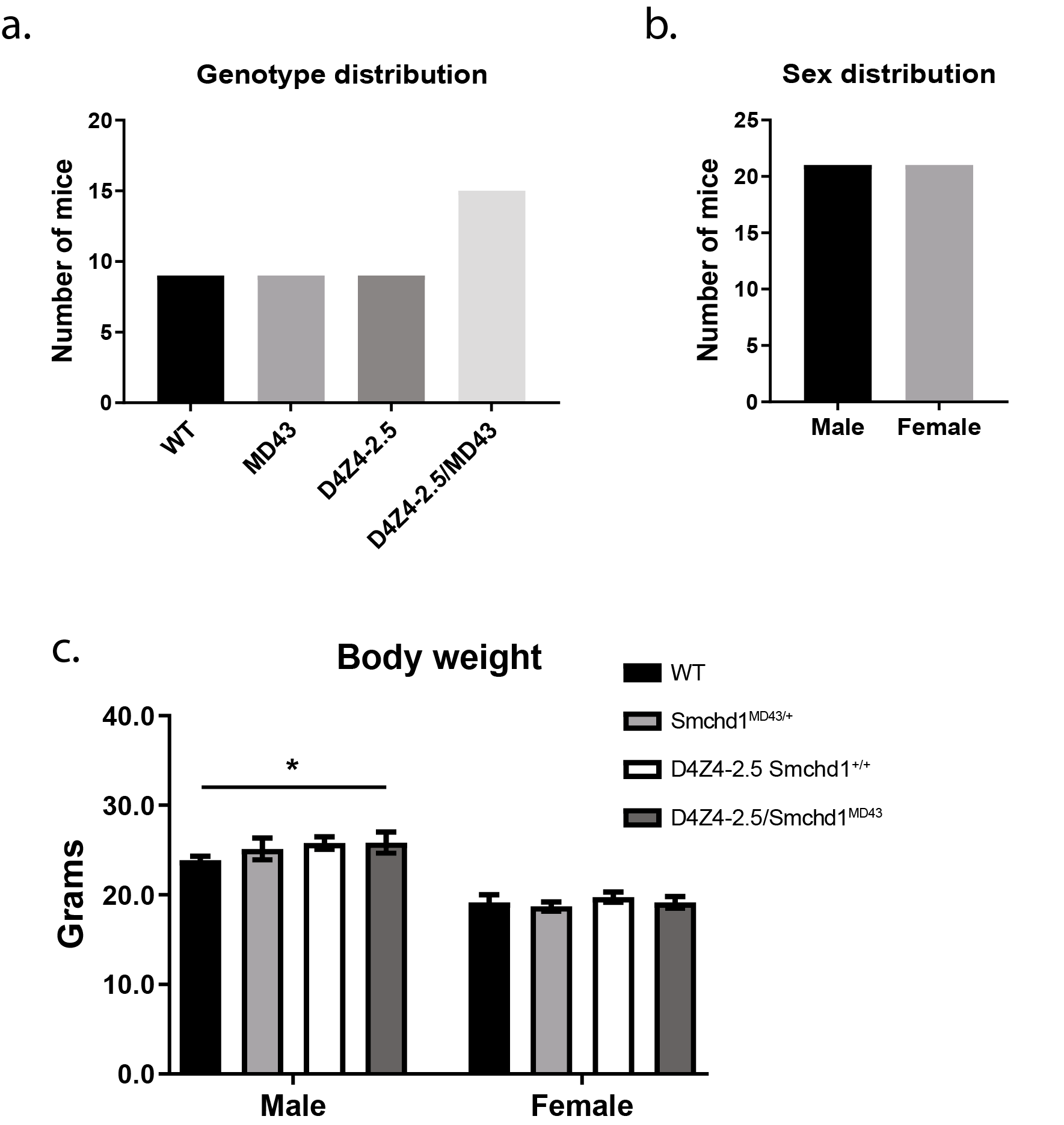

**Supplementary Figure 6:** a. Genotype distribution was not disturbed as tested by a Pearson’s chi-squared test. b. An equal number of male (N=21) and female (N=21) mice were born. c. Body weight (in grams) of male and female mice at two months of age. Bars represent the average body weight per genotype; error bars denote the standard deviation. Statistical analysis was performed by two-way ANOVA followed by a Sidak’s test for post hoc analysis. WT = wild-type; MD43 = *Smchd1^MommeD43/+^*. *P < 0.05.

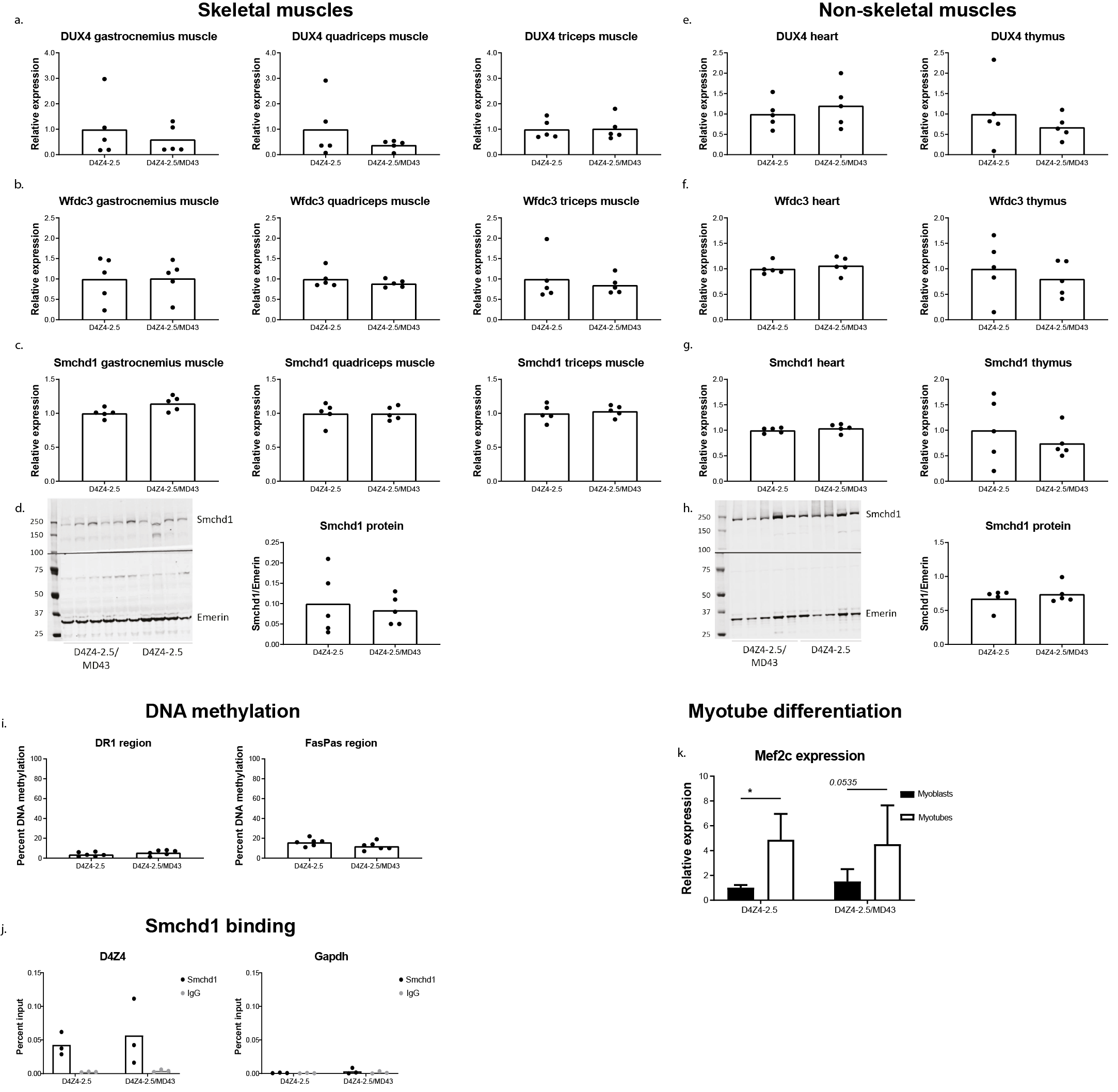

**Supplementary Figure 7:** a-c. a) Relative DUX4 transcript levels, b) Wfdc3 transcript levels, and c) *Smchd1* transcript levels in three different skeletal muscles (gastrocnemius, quadriceps, triceps). d. SMCHD1 protein levels in tibialis anterior muscle. EMERIN was used as a loading control. e-g. e) Relative DUX4 transcript levels, f) Wfdc3 transcript levels, and g) Smchd1 transcript levels in heart and thymus. h. SMCHD1 protein levels in spleen. EMERIN was used as a loading control. i. The average DNA methylation level of 19 CpG dinucleotides within the D4Z4 repeat (the DR1 region) and of 10 CpG dinucleotides just distal to the D4Z4 repeat (the FasPas region) in tail DNA. j. SMCHD1 binding at the *D4Z4* repeat and the *Gapdh* locus in fibroblast cultures. Bars represent the average DNA methylation levels or the SMCHD1 enrichment levels per genotype; each dot represents a single mouse. k. Relative *Mef2c* expression in myoblast and myotube cultures. Bars represent the average transcript level per genotype (average value in D4Z4-2.5 myoblasts is set as 1); error bars denote the standard deviation. Statistical analysis was performed using a two-way ANOVA followed by a Sidak’s test for post hoc analysis. *p < 0.05. Bars represent the average transcript/protein levels per genotype (average value in D4Z4-2.5 muscle is set as 1); each dot represents a single mouse. Statistical analysis was performed using a Student’s t-test except in k. MD43 = *Smchd1^MommeD43/+^*.

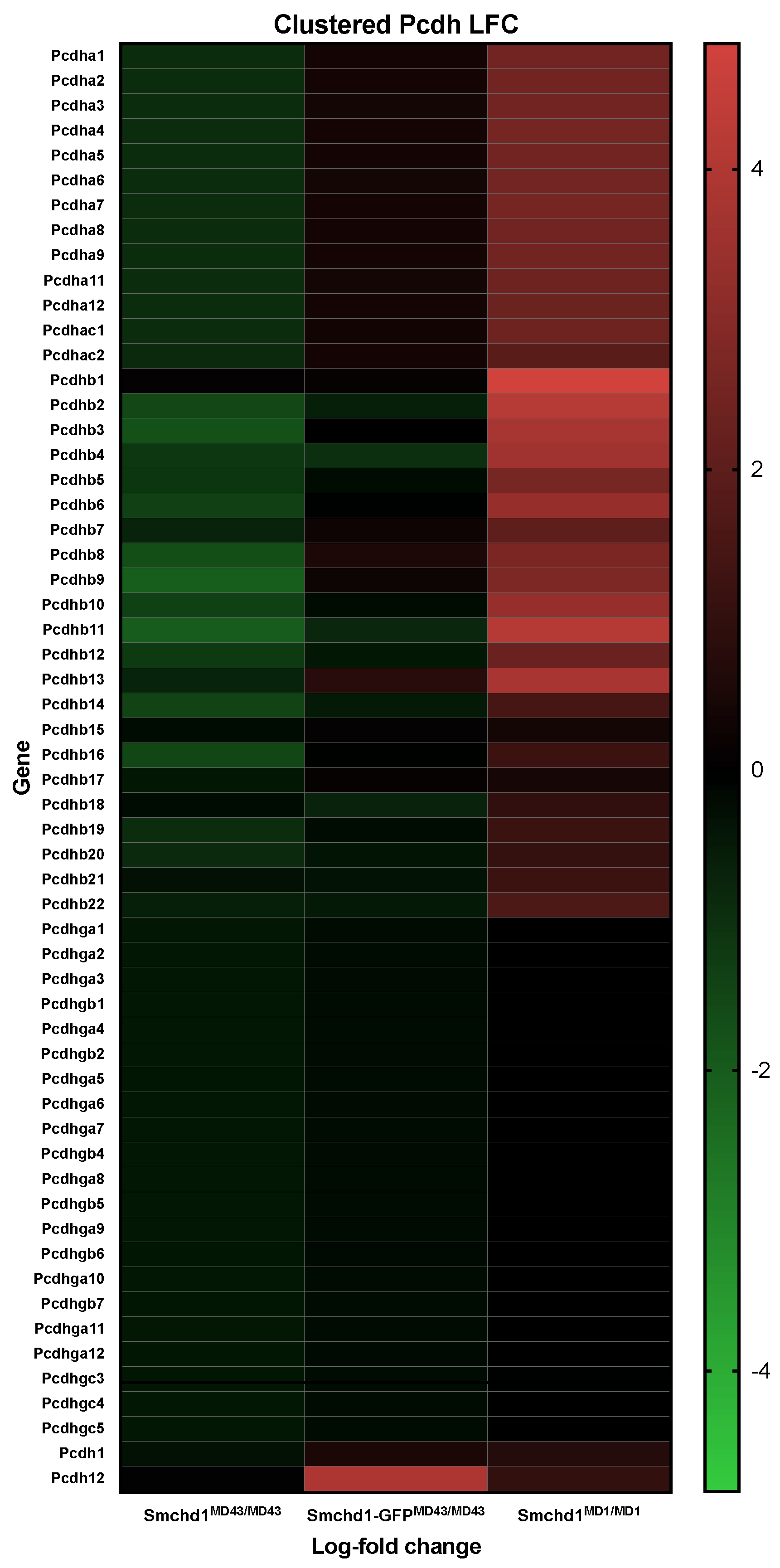

**Supplementary Figure 8:** Heatmap of log-fold change of clustered protocadherin genes in murine chromosome 18. The data is log2 normalized RPKM expression from mRNA sequencing data in *Smchd1^MommeD43/MommeD43^*, *Smchd1-GFP^MommeD43/MommeD43^* and *Smchd1^MommeD1/MommeD1^* NSCs versus their independent controls *Smchd1^+/+^, Smchd1-GFP^+/+^* and *Smchd1^+/+^*, respectively. *MommeD43* abbreviated to *MD43*.

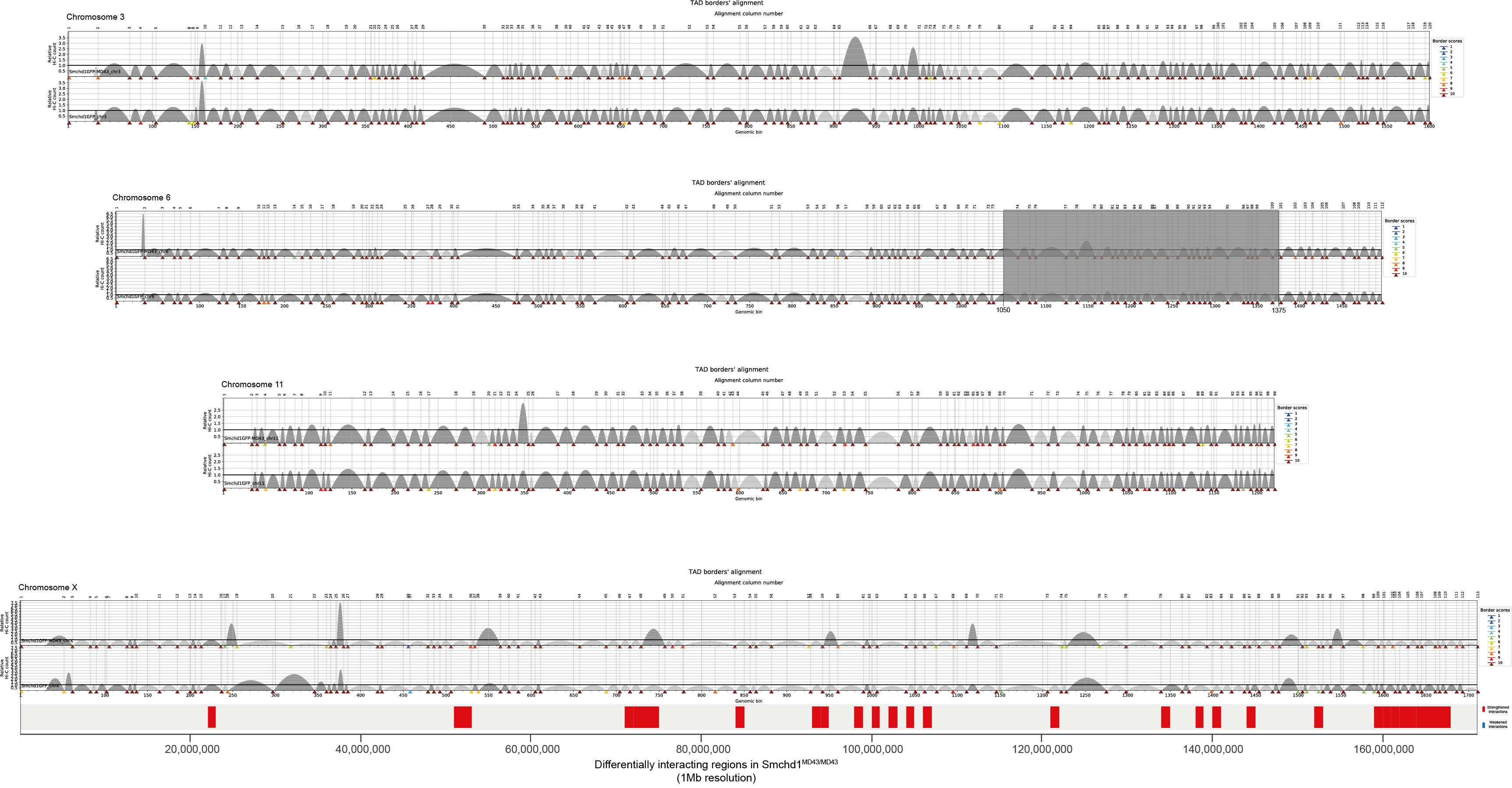

**Supplementary Figure 9.** TAD alignment diagrams from TADbit for chromosomes 3, 6, 11 and X at 100kb resolution. Each diagram shows an individual chromosome with TADs calculated from 3 merged biological replicates for *Smchd1-GFP^MommeD43/GFP-MommeD43^* top) and *Smchd1-GFP^+/+^* NSCs (bottom). TADs with relative Hi-C count >1 are in dark grey. The arrows on the X-axis represent TAD borders, their colour represents their statistical significance (border scores with 10 being highest significance). The Chr6:105Mb-137.5Mb region is greyed out as it was excluded from all analyses because of a chromosomal rearrangement present in the Smchd1-GFP^MD43/MD43^ mouse colony. The units of the X-axis are genomic bins equal to the resolution (100kb). In addition, below the X chromosome’s TAD alignment there is a track showing the statistically significant differentially interacting regions found in HiC data at 1Mb resolution in Smchd1-GFP^MD43/MD43^ compared to Smchd1-GFP^+/+^ NSCs (31 differential interactions FDR<0.1, all strengthened in *Smchd1-GFP^MommeD43-MommeD43^*). *MommeD43* abbreviated to *MD43*.

**Supplementary Tables:**

**Supplementary table 1**

**Data collection and scattering parameters for SAXS analysis**

| **Data collection parameters** | |  | | | |
| --- | --- | --- | --- | --- | --- |
| Instrument |  | | Australian Synchrotron SAXS/WAXS beamline | | |
| Beam geometry |  | | 120 μm point source | | |
| Beam wavelength (Å) |  | | 1.033 | | |
| *q* range (Å^-1^) |  | | 0.0114-0.4 | | |
| Exposure time (seconds) |  | | 2 | | |
| Protein concentration |  | | ~5 mg/ml sample injected via in-line size exclusion chromatography | | |
| Temperature (°C) |  | | 16 | | |
| **Structural parameters** |  | |  | | |
| Protein sample | | | | SMCHD1 111-702 WT | SMCHD1 111-702 A667E |
| *I*(0)(cm^-1^)  [from Guinier] | | | | 0.01806  ±0.00013 | 0.01196  ±0.00012 |
| *Rg* (Å)  [from Guinier] | | | | 31.2  ±0.344 | 31.7  ±0.484 |
| *I*(0)(cm^-1^)  [from P(r)] | | | | 0.01825  ±0.00009 | 0.01198  ±0.00008 |
| *Rg* (Å)  [from P(r)] | | | | 32.17  ±0.206 | 32.21  ±0.273 |
| *D_max_* (Å) | | | | 105 | 105 |
| **Software** |  | |  | | |
| Primary data reduction |  | | ScatterBrain (Australia Synchrotron) | | |
| Data processing |  | | PRIMUS, GNOM | | |

**Summary of melting temperature (Tm) and dissociation constant (*K*_d_) of wild-type and mutants of SMCHD1 hinge**

| **Property of SMCHD1 hinge** | | **Tm (°C) *K*_d_ (*μ*M)** | |
| --- | --- | --- | --- |
| WT | 53.0±1.0 | | 1.5±0.1 |
| R1867G | 42.4±0.1 | | 6.2±0.7 |
| **Dimer Interface mutants** |  | |  |
| D1749A | 45.9±0.3 | | ND. |
| R1762A | 46.8±0.4 | | 3.2±0.2 |
| Y1765A | 45.2±0.4 | | 1.1±0.1 |
| V1774G | 35.0±0.3 | | ND. |
| D1842A | 51.7±0.5 | | 1.2±0.1 |
| R1848A | 42.5±0.1 | | 4.3±1.4 |
| G1872A | 45.9±0.3 | | 1.0±0.1 |
| G1872A, G1875A, G1876A | 38.2±1.2 | |  |
| K1873A* | 67.9±0.3 | | 8.1±1.2 |
| F1874A | 41.4±0.3 | | 1.5±0.1 |
| **Cluster 1 Mutants** |  | |  |
| K1718A | 51.3±0.6 | | 1.4±0.1 |
| R1719A | 52.0±0.8 | | 1.8±0.1 |
| R1771A | 52.7±1.0 | | 2.4±0.2 |
| **Cluster 2 Mutants** |  | |  |
| K1789A | 51.7±0.4 | | 2.3±0.1 |
| R1790A | 54.8±1.1 | | 7.6±0.8 |
| R1796A | 53.4±1.2 | | 4.3±0.3 |
| K1799A | 51.8±0.4 | | 3.4±0.3 |
| **Cluster 3 Mutants** |  | |  |
| R1869A | 61.7±0.3 | | 23.6±4.6 |
| K1873A * | 67.9±0.3 | | 8.1±1.2 |
| K1880A | 44.7±0.2 | | 1.4±0.1 |

K1873A is marked by asterisk as it is categorized as both dimer interface mutant and Cluster 3 mutant**.**

**Supplementary Table 8. Oligonucleotide sequences**

| **Oligo name** | **Purpose** | **Sequence** |
| --- | --- | --- |
| D4Z4 gen F | D4Z4 transgene genotyping | 5’- TGGTCCGTGAAGACATGTGT-3’ |
| D4Z4 gen R | D4Z4 transgene genotyping | 5’- GAGCCCTCAGAGAAGTGGG-3’ |
| Smchd1 ex 15F | MommeD43 genotyping | 5’-TTGTCTTATTTGCAATGTTGGTG-3’ |
| Smchd1 ex 15R | MommeD43 genotyping | 5’-GCTACAGTGCCTAGCCCAAT-3’ |
| MommeD43 probe | MommeD43 genotyping | 5’-AACTGTGCCCATTGCAAAGCTGGATAGGA-3’ |
| D4Z4 ChIP F | D4Z4 ChIP-qPCR | 5’- CCGCGTCCGTCCGTGAAA-3 |
| D4Z4 ChIP R | D4Z4 ChIP-qPCR | 5’-TCCGTCGCCGTCCTCGTC-3’ |
| Gapdh ChIP F | Gapdh ChIP-qPCR | 5’-TGAGCCTCCTCCAATTCAAC-3’ |
| Gapdh ChIP R | Gapdh ChIP-qPCR | 5’-CCAGGAAGACGCTTGAAAAG-3’ |
| DR1 meth F | D4Z4 DR1 methylation | 5’-GGGTTGAGGGTTGGGTTTATA-3’ |
| DR1 meth R | D4Z4 DR1 methylation | 5’- ACAAAACTCAACCTAAAAATATAC -3’ |
| Faspas meth F | D4Z4 Faspas methylation | 5'-ATAGGGAGGGGGTATTTTA-3’ |
| Faspas meth R | D4Z4 Faspas methylation | 5'-ACRATCAAAAACATACCTCTATCTA-3’ |
| Otc F | X chromosome genotyping | 5’-GTTCTTTCGTTTTCCCCTCTC-3’ |
| Otc R | X chromosome genotyping | 5’-GGCATTATCTAAGGAGAAGCATC-3’ |
| Zfy F | Y chromosome genotyping | 5’-GACTAGACATGTCTTAACATCTGTCC-3’ |
| Zfy R | Y chromosome genotyping | 5’-CCTATTGCATGGACAGCAGCTTATG-3’ |
| Smchd1 cKO F2 | Smchd1 fl and del genotyping | 5’-TCAGGTGGTCTCGAGCCC-3’ |
| Smchd1 cKO F4 | Smchd1 fl and del genotyping | 5’-CCATGAGAAGCAATGTGGGA-3’ |
| Smchd1 cKO R1 | Smchd1 fl and del genotyping | 5’-GGACAGCCAAAGTGACACAG-3’ |
| Smchd1 GFP F1 | Smchd1-GFP genotyping | 5’-GCCTGCCTTGCTTCTATGTC-3’ |
| Smchd1 GFP R1 | Smchd1-GFP genotyping | 5’-GCCCCATGAGATTCTGAAAG-3’ |
| Smchd1 GFP R2 | Smchd1-GFP genotyping | 5’-GAATTCAGGGTCAGCTTGC-3’ |
| Momme D1 F | Allelic discrimination for genotyping of Smchd1 | 5’-TCCTCCTTGTGGCCTTGTGG-3’ |
| MommeD1 R | Allelic discrimination for genotyping of Smchd1 | 5’-CGTATTTAAAGGTCCAGCTGTTGC-3’ |
| Taqman Smchd1 wild-type probe used with MommeD1 | Allelic discrimination for genotyping of Smchd1 | VIC-CAGCTTTGGTTGTGCTGT-MGBNFQ |
| Taqman Smchd1 MommeD1 probe | Allelic discrimination for genotyping of Smchd1 | 6FAM-CAGCTTTGGTTATGCTGT-MGBNFQ |
| Taqman Smchd1 wild-type used with MommeD43 probe | Allelic discrimination for genotyping of Smchd1 | VIC-TCCTATCCAGCTTTGCAAT-MGBNFQ |
| Taqman Smchd1 MommeD43 probe | Allelic discrimination for genotyping of Smchd1 | 6FAM-CCTATCCAGCTTTTCAAT-MGBNFQ |
| MommeD43 F | Allelic discrimination for genotyping of Smchd1 | 5’-CCCCAGGCACTGTATGATGAAATAA-3’ |
| MommeD43 R | Allelic discrimination for genotyping of Smchd1 | 5’-CATATTTCCTAATTGTTTTCTCGGCTACTG-3’ |
| sgRNA MommeD43 | gRNA for CRISPR/Cas9 | 5’-TTGCAAAGCTGGATAGGACA-3’ |
| Smchd1 exon 15 MommeD43 | Oligo donor repair template for introducing the MommeD43 mutation by CRISPR/Cas9 | 5’-tgtaaatttttattcagGAACCCCAGGCACTGTATGATGAAATAAAAACTGTGCCTATTGAAAAGCTGGATAGGACAGTAGCCGAGAAAACAATTAGGAAATATGTAGAAGATGAAATGGC-3’ |
